# Ratio-Free Detection and Partial Field Illumination Improve Time-Domain Dynamic Full-Field Optical Coherence Tomography Sensitivity for Retinal Organoid Imaging

**DOI:** 10.64898/2026.04.18.719402

**Authors:** Tual Monfort

## Abstract

Time domain Dynamic full-field optical coherence tomography (D-FFOCT) is a powerful label-free imaging modality that enables functional visualization of cellular activity in living tissues with subcellular resolution. However, its sensitivity remains a major limitation for imaging highly scattering three-dimensional (3D) biological models such as retinal organoids, where incoherent background and inefficient optical flux distribution reduce dynamic contrast and limit imaging depth. In this work, we introduce a ratio-free optical configuration for time-domain D-FFOCT that enables continuous tuning of the sample-to-reference field ratio while minimizing photon losses and suppressing parasitic reflections. This polarization-based architecture allows optimal redistribution of optical flux according to sample scattering conditions and improves sensitivity under both power-limited and dose-limited conditions. Compared with conventional non-polarizing beam splitter configurations, the proposed approach provides a 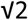-fold (3 dB) sensitivity improvement through optical optimization alone. In addition, we investigate for the first time the use of partial field illumination (PFI) in time-domain D-FFOCT to reduce incoherent background arising from multiple scattering. In retinal organoids imaged at 120 µm depth, PFI yields up to a 14.5-fold (23.2 dB) increase in dynamic signal sensitivity, while preserving functional contrast. When combined, ratio-free detection and PFI provide a cumulative sensitivity improvement of 20.5-fold (26.2 dB). These gains enable improved visualization of photoreceptor precursor organization, rosette structures, and Müller glial cell dynamics in both 3D retinal organoids and 2D cell cultures. This work establishes a practical framework for sensitivity optimization in D-FFOCT and expands its potential for functional imaging, disease modelling, and live-cell monitoring in complex biological systems.

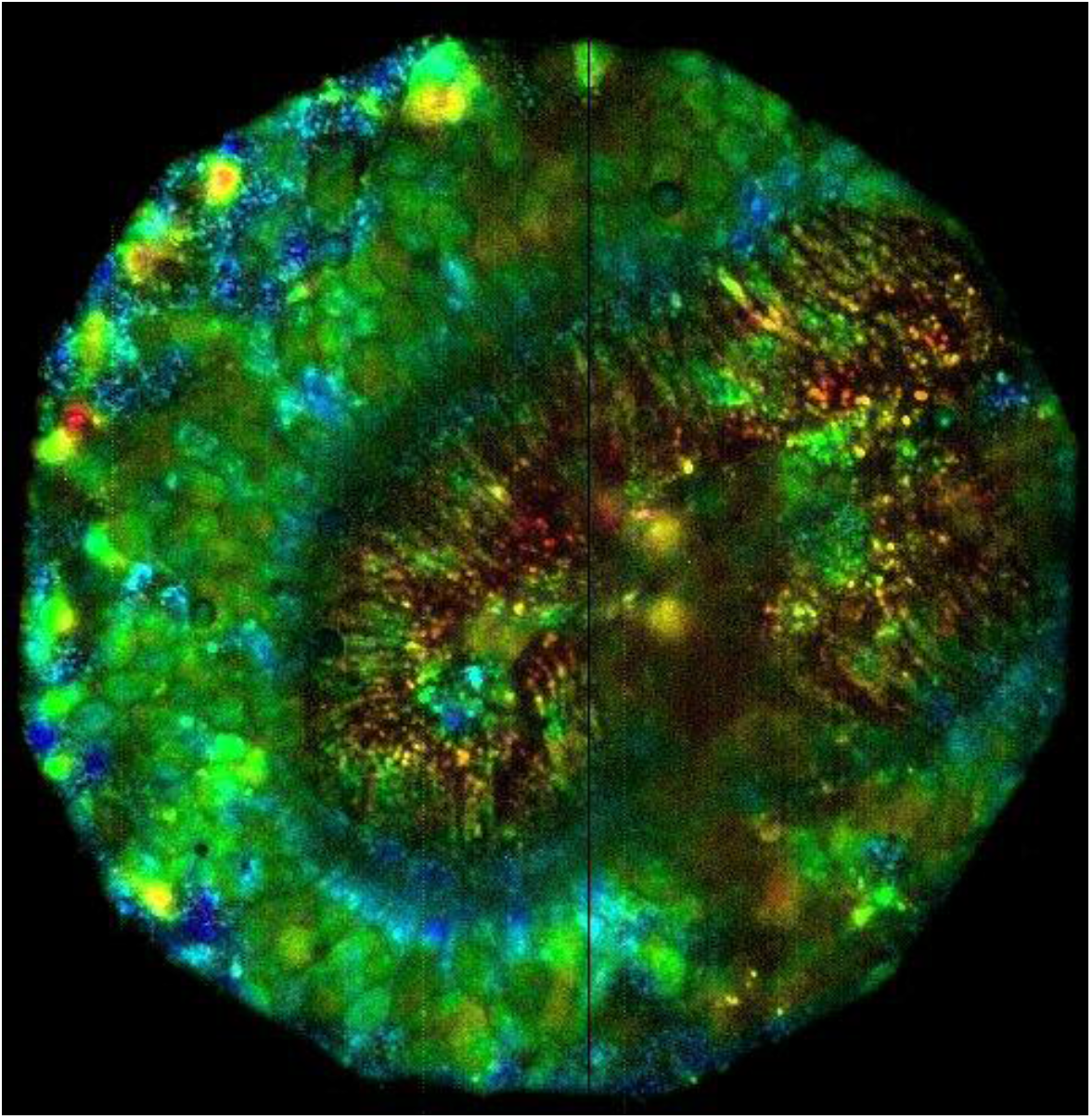

## Introduction

Three-dimensional (3D) cell cultures, organoids, and tissue explants have become essential models for studying human development, disease mechanisms, and therapeutic responses. Compared with conventional two-dimensional (2D) cultures, these systems better reproduce the structural organization, cellular heterogeneity, and microenvironment of native tissues, making them highly relevant for drug screening, disease modelling, and reducing reliance on animal experimentation [1–8]. In ophthalmology, retinal organoids derived from human induced pluripotent stem cells have emerged as particularly valuable models for investigating retinal development, neurodegeneration, inherited retinal diseases, and photoreceptor physiology.

Despite their biological relevance, imaging complex 3D models in a label-free configuration remains challenging. Their thickness often exceeds many times the light transport mean free path, resulting in strong multiple scattering, reduced penetration depth, and poor image contrast compared with 2D cultures [9]. Conventional fluorescence microscopy can provide high specificity, but often requires exogenous markers that may alter cellular physiology, limit long-term observations, or become incompatible with clinical translation. Label-free imaging approaches are therefore highly desirable for continuous live-cell monitoring and functional characterization of biological samples.

Beyond structural imaging, recent advances in live-cell microscopy have highlighted the importance of measuring temporal cellular dynamics as functional biomarkers of tissue physiology [10–21]. Processes such as proliferation, migration, differentiation, apoptosis, and intracellular transport reflect metabolic activity and cellular health, and their dysregulation is associated with numerous pathological conditions including cancer, autoimmune diseases, neurodegeneration, and chronic inflammation. Functional imaging of these dynamic processes is therefore critical for understanding disease progression and evaluating therapeutic responses in physiologically relevant models.

Optical coherence tomography (OCT), and particularly time-domain full-field OCT (FFOCT), addresses many of these challenges by providing label-free, high-resolution en face imaging with submicron axial and lateral resolution [22–26]. By exploiting interference between backscattered light from the sample and a reference field, FFOCT enables optical sectioning, signal amplification, and phase-sensitive detection without mechanical scanning. This configuration is especially well suited for high numerical aperture objectives and has demonstrated cellular-scale imaging in both biological tissues and retinal models.

More recently, dynamic OCT and time domain dynamic full-field OCT (D-FFOCT) have extended this approach from purely structural imaging toward functional imaging of living systems [27, 28]. Rather than relying solely on static reflectivity contrast, D-FFOCT analyzes temporal fluctuations of the interferometric signal to reveal intracellular motion and metabolic activity within dense biological environments. This dynamic contrast has been shown to correlate with cellular activity and can provide a degree of cell-type specificity, offering complementary information to conventional structural FFOCT [9, 24, 29]. D-FFOCT is therefore particularly promising for longitudinal monitoring of organoids, explants, and live cell cultures.

However, sensitivity remains one of the major limitations of D-FFOCT, particularly for highly scattering 3D samples. Current systems typically rely on a 50/50 non-polarizing beam splitter (NPBS), which inherently wastes approximately 75% of the available optical power, including half of the useful OCT signal [9, 28, 30, 31]. In addition, incoherent parasitic reflections from cube beam splitters and multiply scattered background light from the sample occupy a significant fraction of the detector dynamic range, directly reducing sensitivity. Although increasing the optical power could, in principle, compensate for these losses, practical limitations arise from the spatially incoherent nature of the illumination and its Köhler coupling to the microscope objective—particularly in long-arm interferometers compatible with commercial systems [9]. In practice, phase stability requires the use of cube beam splitters, which introduce non-negligible incoherent parasitic reflections. These contributions occupy a fraction of the detector dynamic range and directly reduce sensitivity. Consequently, increasing the source power or altering the splitting ratio also amplifies these parasitic signals, thereby limiting the net gain achievable through power scaling alone.

Furthermore, the reference field is commonly attenuated using neutral density filters or partially reflective mirrors to balance the interferometric signal with the incoherent background [9, 28, 30, 31]. While effective under specific conditions, this strategy introduces substantial photon loss and limits adaptability across different imaging depths and sample scattering conditions. Alternative light-efficient OCT configurations have been proposed [30, 32, 33], but they are not directly compatible with standard high-numerical-aperture FFOCT implementations or rely on different OCT modalities.

In this work, we revisit a polarization-based optical configuration originally introduced by Beaurepaire et al. [23] and apply it to D-FFOCT for the first time. This ratio-free architecture enables continuous tuning of the sample-to-reference field ratio while minimizing photon losses and strongly suppressing parasitic reflections. In parallel, we investigate the impact of partial field illumination (PFI) [34] in D-FFOCT to reduce incoherent background arising from multiple scattering. Together, these two approaches provide a practical strategy for sensitivity optimization under both power-limited and dose-limited conditions.

We demonstrate that ratio-free detection improves sensitivity by 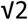 (3 dB) compared with conventional NPBS configurations, while partial field illumination provides up to a 14.5-fold (23.2 dB) increase in dynamic signal sensitivity in retinal organoids at 120 µm depth. Combined, these approaches yield a cumulative improvement of 20.5-fold (26.2 dB), enabling enhanced visualization of photoreceptor precursor organization, rosette structures, and Müller glial cell dynamics. These results establish a new framework for improving D-FFOCT performance and expand its potential for functional imaging, disease modelling, and live-cell monitoring in complex biological systems.

## Results

### 1. Signal model and sensitivity in D-FFOCT

In time-domain FFOCT, the signal detected on a single camera pixel arises from the superposition of coherent and incoherent contributions. Assuming that the depth of field is smaller than the coherence volume, the detected number of photoelectrons per pixel per frame can be expressed as:

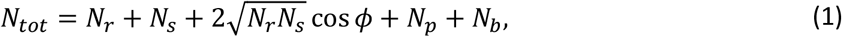

where *N*_*r*_, and *N*_*s*_ denote the detected reference and coherent sample photons, respectively, *N*_*p*_ represents parasitic incoherent reflections originating from optical components, and *N*_*b*_ corresponds to the incoherent background generated by multiple scattering within the sample. The phase difference between the reference and sample fields is represented by *ϕ*.

In practical operation, the camera is used close to its full-well capacity *N*_*FWC*_ in order to operate in a shot-noise-limited regime where read noise becomes negligible. Under these conditions:

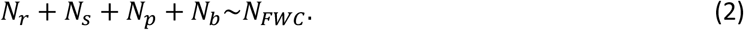

The relevant quantity is therefore not the absolute interferometric signal itself, but the coherent modulation relative to the detector DC level.

In dynamic FFOCT (D-FFOCT), the useful contrast originates from temporal fluctuations of the coherent interference term. For a time series of *M* statistically independent frames, and independently of the initial phase of the scatterers [26], the signal-to-noise ratio (SNR) can be expressed as:

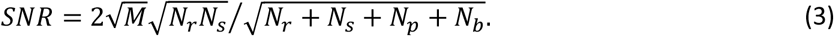

In the biologically relevant regime where the coherent sample signal remains much smaller than the reference and incoherent background contributions (*N*_*r*_+*N*_*p*_+*N*_*b*_*≫ N*_*s*_), the normalized sensitivity becomes:

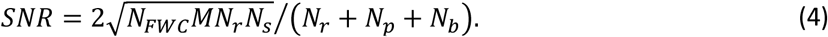

This expression highlights the central limitation of D-FFOCT sensitivity: the coherent signal is not only constrained by the amount of useful sample photons, but also by the total incoherent DC contribution filling the detector dynamic range. In highly scattering biological samples such as retinal organoids, the multiple-scattering background and parasitic reflections can dominate the detected flux, substantially reducing dynamic contrast.

This sensitivity bottleneck is particularly critical for deep imaging, where weak coherent signals must be detected in the presence of strong incoherent background. Optimizing the distribution of optical flux between the sample and reference arms therefore becomes essential for improving D-FFOCT performance.

### 2. Optical flux optimization: ratio-free D-FFOCT

#### 2.1 Conventional NPBS configuration

In conventional D-FFOCT systems, the interferometer is based on a 50/50 non-polarizing beam splitter (NPBS), which imposes a fixed optical power distribution between the sample and reference arms. For a reference reflectivity 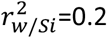 9, the coherent sample and reference photon counts can be written as:

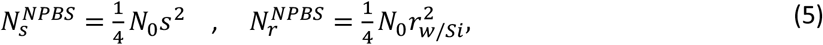

where *N*_0_ is the number of photons incident on the interferometer during one exposure, and *s* and *r*_*w/si*_ represent the field reflectivities of the sample and reference, respectively. The quantum efficiency is omitted for simplification purposes.

The incoherent contribution generated by multiply scattered light within the sample is modeled as:

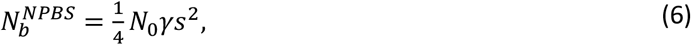

where *γ* is the ratio between incoherent and coherent sample return, typically *γ ≫*1 in highly scattering media. In addition, the NPBS introduces parasitic optical incoherent reflections

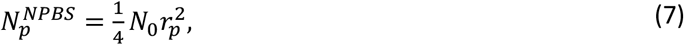

where 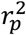 represents the effective optical parasitic reflection term.

Substituting these contributions into the SNR expression yields the reference sensitivity:

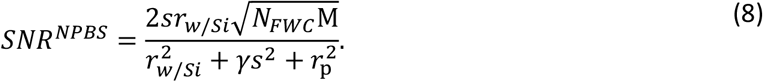

This formulation shows that sensitivity is fundamentally limited by three competing contributions: the reference intensity, the incoherent sample background, and parasitic reflections from the optical system. Because the 50/50 beam splitter imposes a fixed ratio between the sample and reference arms, no adaptation is possible when sample scattering conditions vary with depth or biological complexity.

As a result, conventional NPBS-based D-FFOCT operates far from the theoretical optimum in highly scattering samples, where dynamic contrast is often constrained by detector saturation rather than by the availability of coherent signal.

### 2.2 Ratio-free polarization architecture

To overcome these limitations, we implemented a polarization-based ratio-free architecture composed of a polarizer (Pol.1), a polarizing beam splitter (PBS), quarter-wave plates in both interferometer arms, and an analyzer (Pol.2), as shown in Fig. 1.

**Figure 1.**
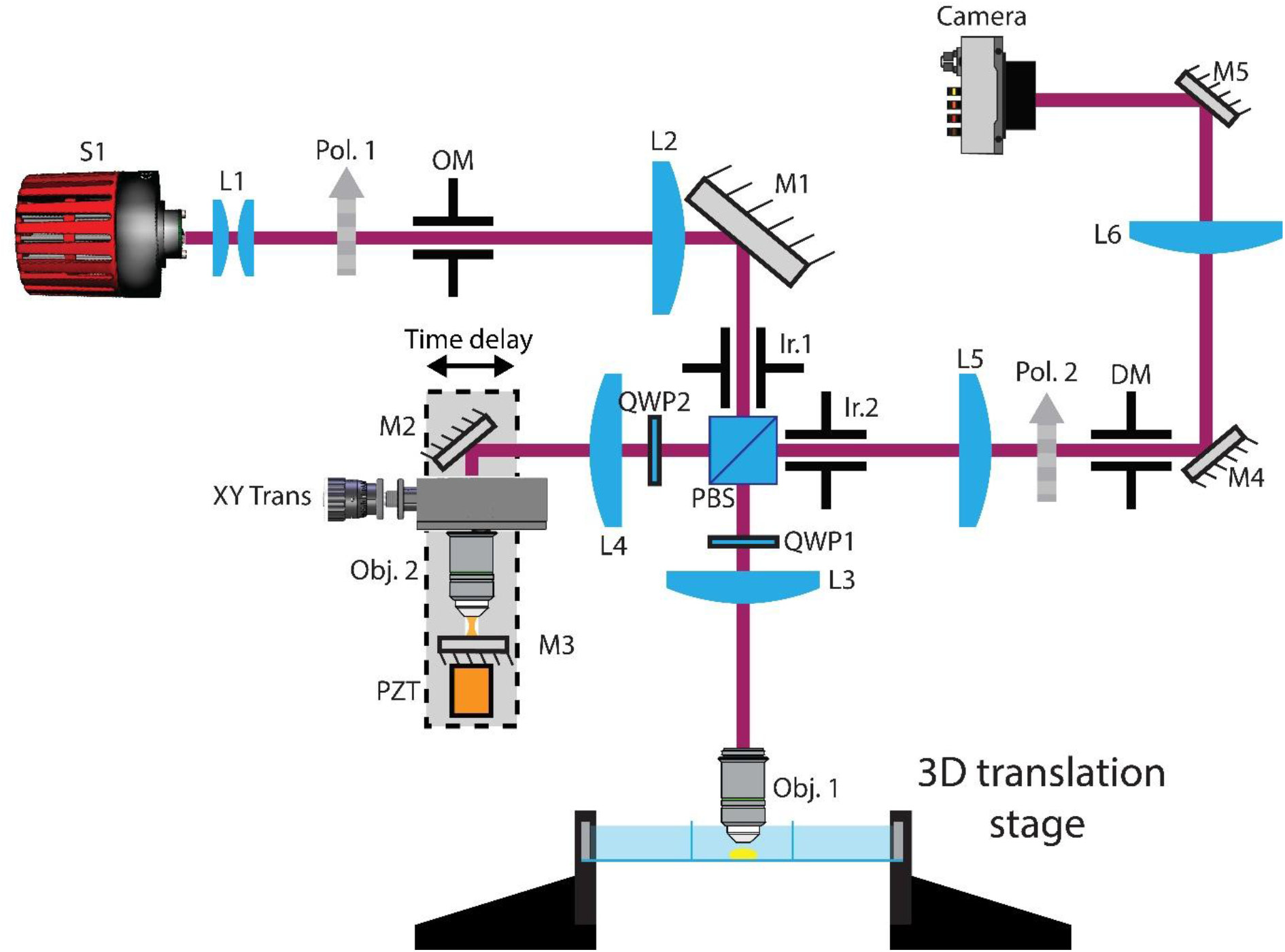
Optical layout of ratio-free dynamic full-field optical coherence tomography (D-FFOCT) system. A mounted light-emitting diode (LED, 810 nm center wavelength, 30 nm bandwidth) was used as an extended spatially incoherent source (S1). A first lens pair (L1) imaged the source onto the illumination mask (OM). A first polarizer (Pol.1), combined with a polarizing beam splitter (PBS), controlled the sample-to-reference optical flux ratio by adjusting the polarization balance between the two interferometer arms. A doublet (L2) relayed the illumination mask to the back focal planes of the microscope objectives (Obj.1 and Obj.2) through the intermediate relay lenses (L3 and L4), forming the Linnik interferometer. The illumination iris (Ir.1), positioned at the focal plane of L2 and conjugated to the sample plane, was used for partial field illumination (PFI) to reduce incoherent background generated by multiple scattering. A piezoelectric actuator (PZT) and a linear delay stage introduced dynamic and static phase shifts, respectively. Quarter-wave plates (QWP1 and QWP2) rotated the polarization state after reflection from either the sample or the reference mirror, enabling polarization routing through the PBS and strong suppression of parasitic reflections. A second polarizer (Pol.2) acted as an analyzer to project the orthogonal sample and reference fields onto a common polarization axis before detection by the cMOS camera. This ratio-free polarization architecture enables continuous tuning of the sample-to-reference field ratio, improves detector dynamic range usage, and provides the optical basis for both sensitivity enhancement and partial field illumination in D-FFOCT.

In this configuration, the angle *α* of Pol.1 controls the distribution of optical power between the sample and reference arms by balancing the horizontal and vertical polarization components of the spatially incoherent source. The analyzer angle *β* then controls the detected projection of the returned fields.

The detected coherent photon counts become:

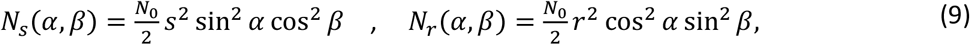

where *r* is the reflectivity of the reference mirror, typically close to unity. The incoherent sample background remains proportional to the coherent sample return:

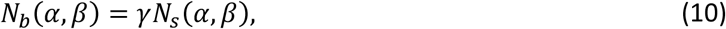

while parasitic optical reflections are strongly suppressed by polarization routing and can be neglected at first order:

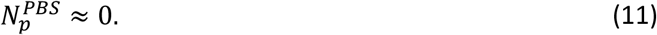

The corresponding interferometric sensitivity becomes:

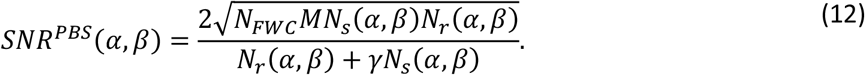

Unlike the conventional NPBS configuration, this ratio-free architecture enables continuous tuning of the sample-to-reference field ratio and direct adaptation to the scattering properties of the biological sample.

Fig. 2a shows the theoretical SNR ratio between the ratio-free PBS architecture and the conventional NPBS configuration. A clear sensitivity advantage is observed across a wide range of configurations, with improvements exceeding 25 dB under ideal optical balancing conditions.

**Figure 2.**
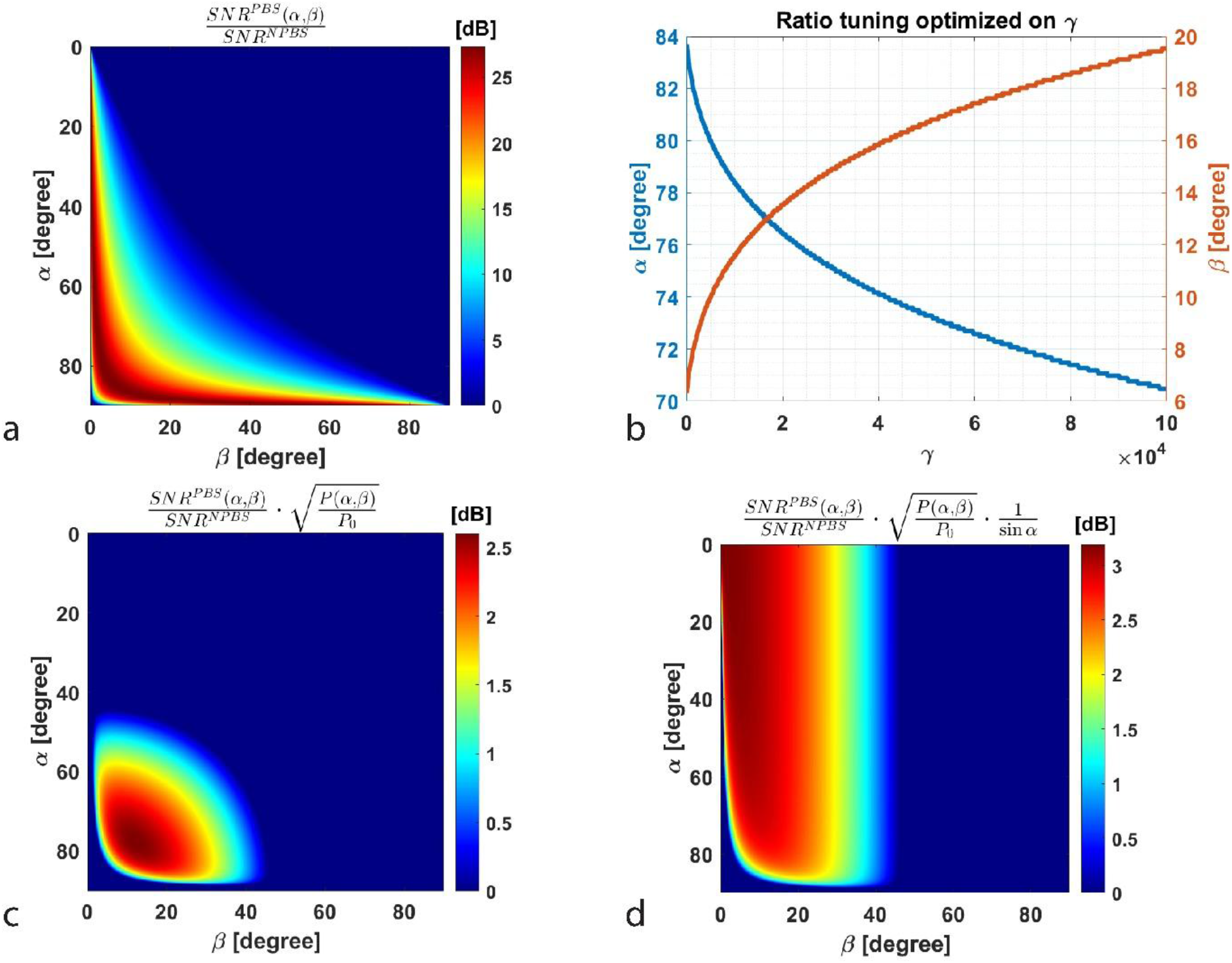
Comparison of signal-to-noise ratio (SNR) between ratio-free PBS and conventional NPBS D-FFOCT configurations. **a** Theoretical SNR ratio between the ratio-free polarization-based architecture (PBS) and the conventional 50/50 non-polarizing beam splitter (NPBS) configuration as a function of the polarizer angles α (Pol.1) and β (Pol.2). The ratio-free configuration provides a substantial sensitivity advantage across a broad range of optical ratios. **b** Dependence of optimal SNR on the selected sample-to-reference ratio under increasing incoherent background contributions ϒ, highlighting the importance of adaptive ratio tuning in highly scattering samples. **c** SNR improvement obtained under fixed available optical power after including the accumulation gain associated with detector full-well saturation. **d** Global experimental sensitivity gain achieved using the ratio-free architecture compared with the conventional NPBS configuration, showing an overall improvement of approximately 3 dB 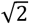-fold. These results demonstrate that sensitivity optimization depends critically on optical flux distribution rather than total illumination power alone.

Importantly, the maximum sensitivity strongly depends on the selected values of *α* and *β*, as shown in Fig. 2b, relative to the amount of incoherent light detected. In highly scattering samples where incoherent background dominates, optimal performance is achieved by reducing the sample illumination fraction while maintaining sufficient reference intensity. This ratio flexibility is particularly important for deep tissue imaging, where multiple scattering progressively fills the detector dynamic range.

### 3. Sensitivity gain mechanisms

#### 3.1 Accumulation gain under full-well constraint

Because both optical configurations use the same detector, the exposure time required to reach camera saturation depends directly on the total detected DC flux. In the shot-noise-limited regime, the saturation time scales inversely with the detected optical power:

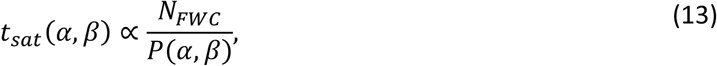

where the total detected DC contribution is given by:

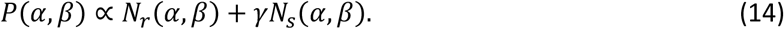

For a fixed total acquisition time, a shorter saturation time allows a larger number of independent frames to be accumulated. Since D-FFOCT sensitivity scales with the square root of the number of frames, this produces an additional accumulation gain:

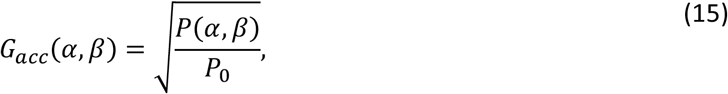

where *P*_0_ corresponds to the DC level of the conventional NPBS configuration. For example, if the ratio-free polarization architecture reaches full-well capacity twice as fast as the conventional NPBS setup, twice as many independent frames can be acquired over the same experimental duration, resulting in an additional sensitivity improvement of 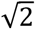.

Fig. 2c shows the contribution of this accumulation term to the global SNR comparison. Under fixed available optical power, the ratio-free configuration provides an additional sensitivity improvement of approximately 2.6 dB compared with the classical NPBS architecture.

#### 3.2 Dose-limited operation

A major advantage of the ratio-free architecture is the decoupling of sample illumination from reference intensity. In the conventional NPBS configuration, increasing the reference field necessarily increases the optical dose delivered to the sample. In contrast, the polarization-based architecture allows these two quantities to be adjusted independently.

Because only a fraction sin^2^ *α* of the incident light is directed toward the sample, the optical dose per frame scales as:

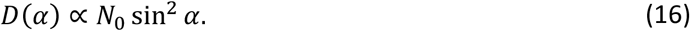

For a fixed maximum allowable sample dose—particularly important for clinical retinal imaging and long-term live-cell observations—the total incident photon flux can therefore be increased as:

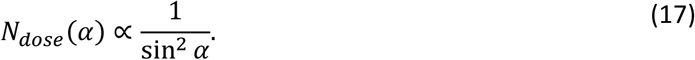

This additional available flux increases the coherent interference term, which scales with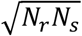, resulting in a dose-limited gain factor:

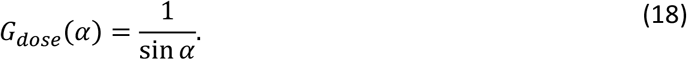

Thus, by reducing the fraction of light directed toward the sample while preserving reference intensity, the ratio-free architecture maximizes the information extracted from a constrained photon budget without increasing biological irradiation.

This feature is particularly relevant for retinal applications, where optical dose must remain strictly below safety limits, such as clinical imaging, as well as for long-term imaging of sensitive biological samples such as organoids and stem-cell-derived cultures.

#### 3.3 Global performance

Combining the intrinsic SNR improvement, the accumulation gain, and the dose-limited operation yields the global merit factor:

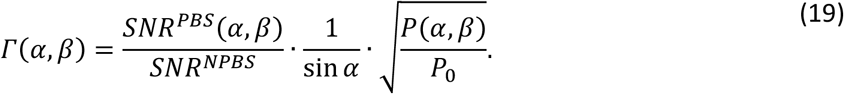

This expression summarizes the full benefit of the ratio-free architecture under realistic experimental conditions.

Experimentally, the ratio-free D-FFOCT configuration produces an overall sensitivity improvement of approximately 3 dB compared with the conventional NPBS setup (Fig. 2d), corresponding to a 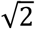-fold gain obtained purely through optical flux optimization.

The need for brighter spatially incoherent sources therefore becomes particularly relevant, as higher source power can be efficiently exploited in this ratio-free configuration, where parasitic reflections are strongly suppressed through polarization rejection.

In highly scattering samples (*γ ≫*1) where multiple scattering occupies a large fraction of the detector dynamic range, this ratio flexibility becomes critical. This is precisely the regime encountered when imaging retinal organoids several transport mean free paths deep, where conventional NPBS-based systems rapidly become background-limited.

Beyond the sensitivity gain itself, this architecture provides a major translational advantage for clinical imaging applications: it maximizes usable interferometric contrast under strict optical safety constraints, making it particularly well suited for retinal imaging and future in vivo applications.

## 4. Biological Validation in Retinal Organoids and Müller Glial Cells

### 4.1 Ratio-Free D-FFOCT Imaging of Retinal Organoids

To validate the practical impact of the ratio-free architecture, we performed D-FFOCT imaging of human retinal organoids at day 150 of differentiation using the optical configuration described in Fig. 1. Imaging was performed at a depth of 50 µm using a water-immersion objective of numerical aperture 0.5.

Despite the relatively moderate numerical aperture, the ratio-free configuration enabled clear visualization of individual photoreceptor precursor boundaries and improved delineation of local cellular organization (Fig. 3a–c). Cellular contours were sufficiently well resolved to allow segmentation of individual cells, demonstrating that the sensitivity improvement obtained through optical flux optimization directly translates into enhanced biological interpretability.

**Figure 3.**
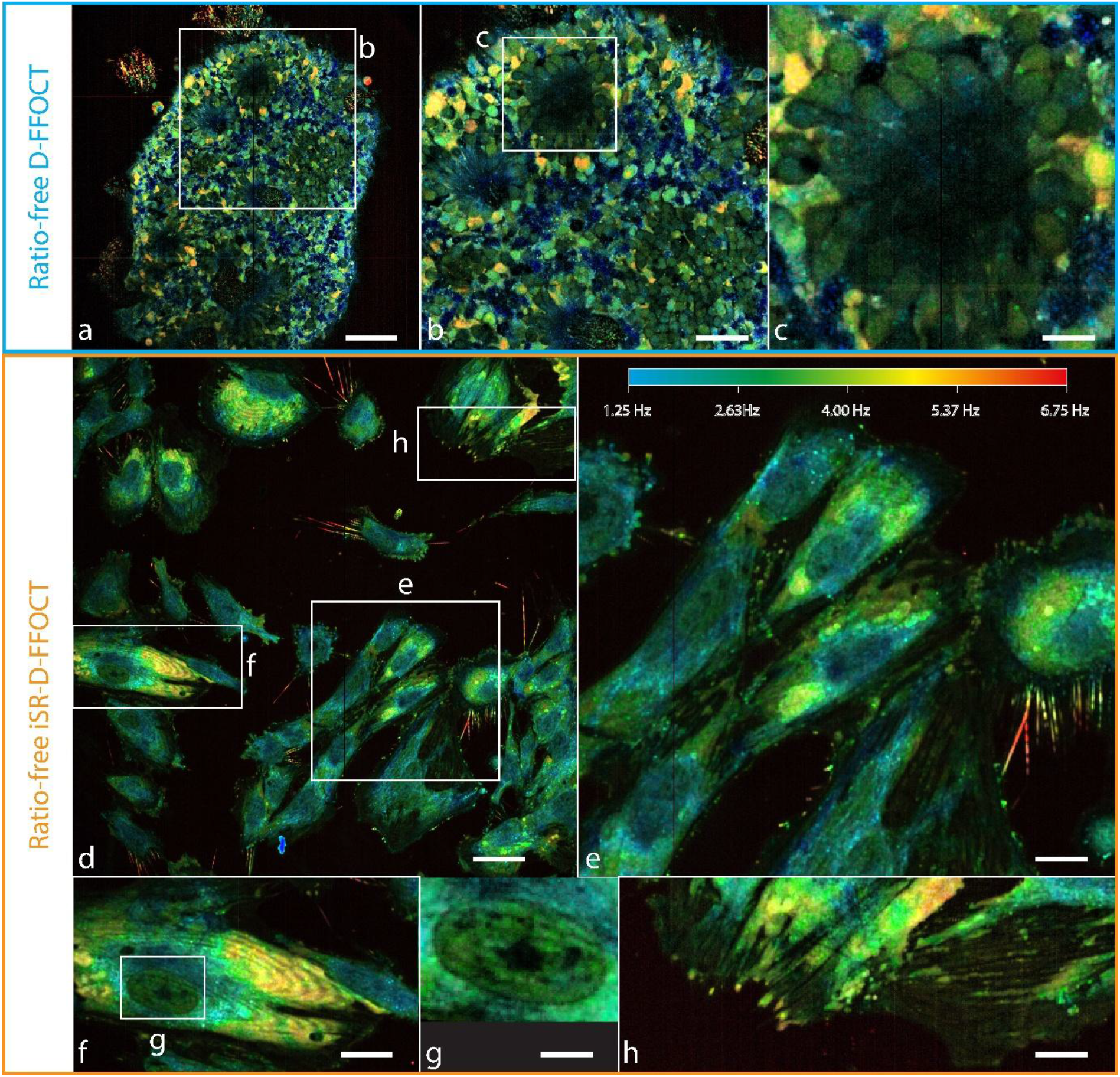
Ratio-free D-FFOCT and interface self-referenced D-FFOCT (iSR-D-FFOCT) imaging of retinal organoids and Müller glial cells. **a** Ratio-free D-FFOCT image of a day-150 retinal organoid acquired at 50 µm depth. **b** Higher magnification of **a**, showing improved visualization of individual photoreceptor precursor boundaries and local tissue organization. **c** Zoom-in of **b** highlighting a retinal rosette structure, a region that is typically difficult to image due to strong multiple scattering and local disruption of retinal lamination. **d** iSR-D-FFOCT image of human induced pluripotent stem cell-derived Müller glial cells cultured in 2D. **e** Higher magnification of **d** showing several Müller cells with apical microvilli-like structures and bright dynamic intracellular compartments suggestive of highly active membrane-associated or metabolic processes. **f** Zoom-in showing interference fringes generated by interference between the plastic–sample interface and the upper membrane– medium interface. **g** Higher magnification highlighting a clearly resolved nucleus with visible contour definition and dark low-dynamic structures consistent with heterochromatin organization. **h** Zoom-in showing highly dynamic protrusive structures resembling filopodia-like extensions. Hue scales linearly from 1.25 to 6.75 Hz in all images. Scale bars: 80 µm in **a**, 40 µm in **b**, 12 µm in **c**, 50 µm in **d**, 22 µm in **e**, 24 µm in **f**, 10 µm in **g**, and 20 µm in **h**.

Particularly notable was the visualization of rosette structures, as highlighted in Fig. 3c. These structures are notoriously difficult to image because they are associated with local disruption of retinal lamination, increased optical aberrations, and strong multiple scattering caused by tissue disorganization. In conventional full-field detection, these regions often appear poorly contrasted or obscured by incoherent background. In contrast, the ratio-free configuration enabled clear structural identification of rosettes and surrounding photoreceptor precursors, supporting deeper functional analysis of retinal organization.

Dead or metabolically inactive cells could also be identified through their characteristic saturated, speckled, and blue-shifted dynamic appearance. In the present experiments, this was partly attributed to imaging performed outside the incubator at ambient temperature, illustrating the sensitivity of D-FFOCT dynamic contrast to physiological state and environmental conditions.

### 4.2 Interface Self-Referenced D-FFOCT of Müller Glial Cells

The ratio-free architecture also enables an optimal self-referenced configuration of D-FFOCT, referred to as interface self-referenced D-FFOCT (iSR-D-FFOCT), obtained by setting *α* = 90degrees and *β* = 0 degrees.

In this configuration, interference occurs between the backscattered sample field and the specular reflection generated at the plastic–sample interface (bottom of the multiwell plate), creating an intrinsically stable OCT reference. This self-referenced approach provides strong interferometric contrast while greatly reducing sensitivity to mechanical vibrations and phase instability [40].

Compared with conventional NPBS-based iSR-D-FFOCT, the polarization implementation preserves twice as much useful optical flux originating from the sample (sample field plus reference field), resulting in an additional 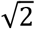-fold (3 dB) sensitivity improvement.

This configuration was applied to 2D cultures of human induced pluripotent stem cell-derived Müller glial cells (Fig. 3d–h). Cellular morphology was visualized with high dynamic contrast, including nuclei, microvilli-like apical structures, and highly active protrusive regions resembling filopodia-like extensions.

Fig. 3g highlights a particularly well-resolved nucleus with visible contour definition and dark low-dynamic structures consistent with heterochromatin organization. Figure 3h reveals proliferative or protrusive structures with increased dynamic activity at the cellular edge, suggesting active membrane remodeling.

Interestingly, large bright dynamic pockets with characteristic yellow hue (~5 Hz) were observed for the first time in 2D Müller cell cultures using iSR-D-FFOCT (Fig. 3e). Their biological origin remains under investigation, but their strong dynamic signature suggests highly active intracellular or membrane-associated processes.

Interference fringes were also observed in some cells (Fig. 3f), resulting from interference between two specular interfaces: the plastic–sample interface and the upper membrane–medium interface. While such fringes can be reduced using broader spectral bandwidth illumination or higher numerical aperture objectives, they also provide a unique opportunity to probe nanometric membrane deformations and subtle 3D morphological variations.

Time-lapse imaging over several hours (Supplementary Video S1) showed limited large-scale cellular displacement, suggesting that the dominant dynamic contrast originated from intracellular activity rather than cell migration. This highlights the ability of iSR-D-FFOCT to probe fine functional dynamics in relatively stable adherent cultures.

Together, these results demonstrate that the ratio-free architecture significantly improves both conventional D-FFOCT and self-referenced D-FFOCT, enabling functional imaging of subtle biological structures that remain difficult to detect using standard configurations.

## 5. Multi-Scattering Mitigation Using Partial Field Illumination

### 5.1 Partial Field Illumination Strategy

Once optical flux distribution was optimized using the ratio-free architecture, we next addressed the second major limitation of D-FFOCT sensitivity: the incoherent background generated by multiple scattering within the sample.

In highly scattering biological tissues such as retinal organoids, multiply scattered photons contribute strongly to the detector DC level without participating in coherent interference. These incoherent contributions occupy a significant fraction of the camera dynamic range and directly reduce the detectability of weak dynamic signals, particularly at depth.

To reduce this background contribution, we implemented partial field illumination (PFI) using the illumination iris (Ir.1 in Fig. 1), thereby restricting the illuminated field of view while preserving interferometric detection. This approach reduces the amount of multiple scattered light reaching the detector at the expense of decreasing the imaging area, effectively improving the coherent-to-incoherent signal ratio.

PFI was evaluated in retinal organoids imaged at a depth of 120 µm, a regime where multiple scattering strongly limits conventional full-field D-FFOCT performance. A total of 512 raw interferometric frames were acquired at 100 Hz for both full-field illumination (100%) and partial field illumination (8%), while maintaining the same reference-to-sample ratio.

Exposure times were adjusted to preserve comparable detector filling conditions: 3.08 ms for full-field illumination and 9.85 ms for 8% PFI.

### 5.2 Static and Dynamic Signal Enhancement

Fig. 4 compares the effect of PFI on both static and dynamic D-FFOCT signals.

**Figure 4.**
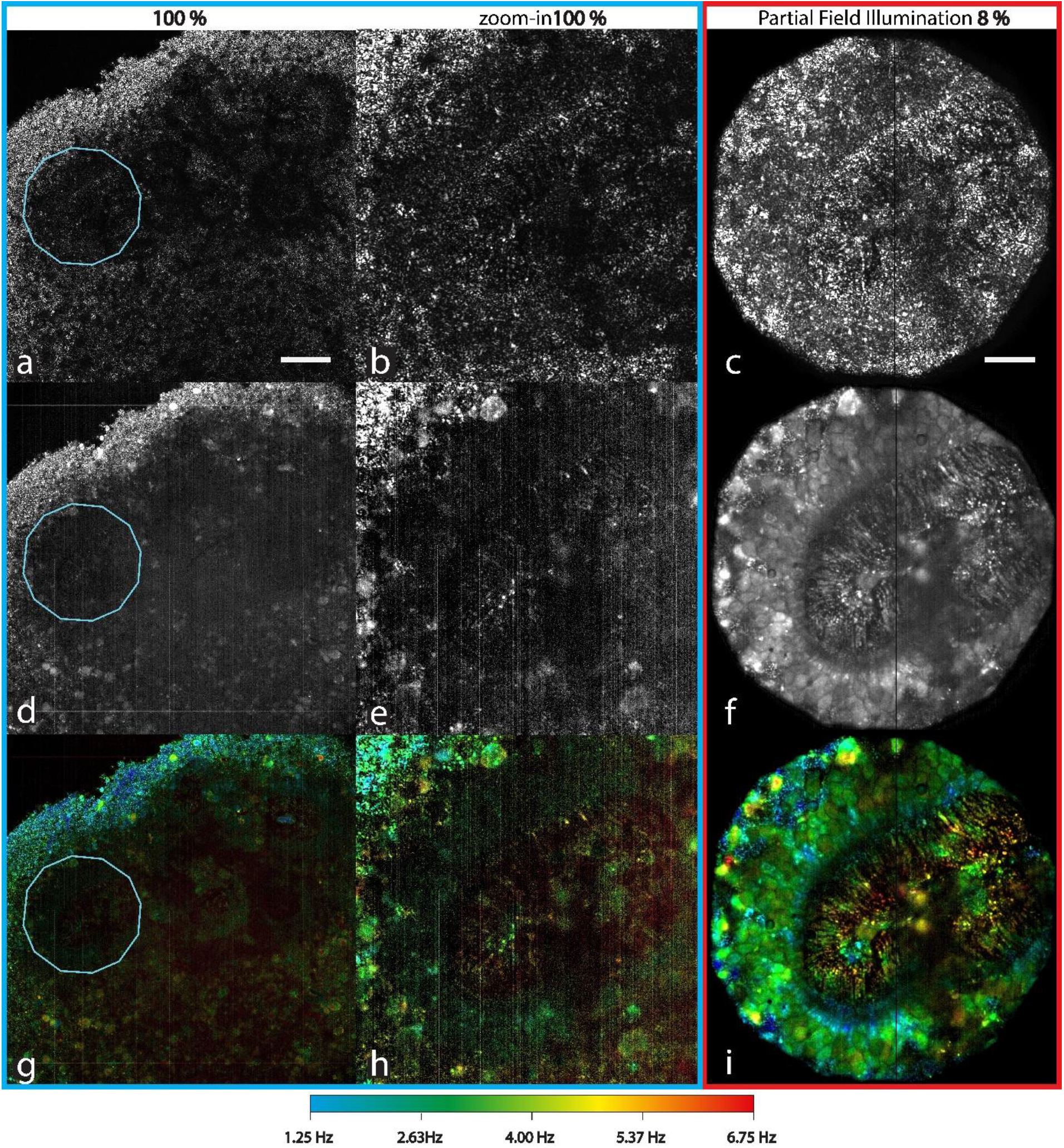
Effect of partial field illumination (PFI) on static and dynamic D-FFOCT sensitivity in retinal organoids. **a–i** Static and dynamic D-FFOCT data acquired from a retinal organoid at 120 µm depth using 512 raw interferometric frames acquired at 100 Hz. Exposure times were 3.08 ms for full-field illumination (100%) and 9.85 ms for partial field illumination (8%), while maintaining a constant reference-to-sample ratio. **a–c** Static two-phase FFOCT signal. **d–f** Phase Fluctuation Index. **g– i** Hue–Saturation–Brightness rendering combining the first two moments of the magnitude spectral density and the Phase Fluctuation Index. **a, b, d, e, g**, and **h** correspond to conventional full-field illumination (100%), whereas **c, f**, and **i** correspond to 8% partial field illumination. The blue ROI indicates the effective illuminated field corresponding to the 8% PFI condition. Partial field illumination substantially improves both structural and dynamic contrast, enabling clear visualization of retinal rosette structures and cellular morphology that remain poorly resolved under conventional illumination. Scale bars: 50 µm in **a–c** and 25 µm in **d–i**. Hue scales linearly from 1.25 Hz (blue) to 6.75 Hz (red).

In the conventional full-field configuration, static structural contrast remained partially visible (Fig. 4a– b), but dynamic contrast metrics such as the Phase Fluctuation Index (PFI metric), mean of the magnitude spectral density (MSD), and standard deviation of MSD showed limited cellular information (Fig. 4d,e,g,h). In particular, rosette structures (one of them is circled by a blue ROI in Fig.4 a,d,g) and individual cellular boundaries remained poorly resolved because the dynamic signal amplitude was close to the system noise floor.

Under 8% partial field illumination, both structural and dynamic contrast improved markedly (Fig. 4c,f,i). Cellular morphology became clearly visible, and rosette organization could be identified with substantially improved contrast. Dynamic cellular activity was also more readily distinguished, revealing structures that remained undetectable under conventional full-field illumination.

This result is particularly important because D-FFOCT relies primarily on weak temporal fluctuations rather than strong static reflectivity. While structural FFOCT may remain interpretable even under moderate incoherent background, dynamic FFOCT requires the useful signal to exceed the camera noise floor. PFI therefore has a proportionally stronger impact on dynamic imaging than on static imaging.

The improved visualization observed in retinal rosettes further supports the hypothesis that scattering, rather than optical aberration alone, is the dominant factor limiting deep imaging performance in these highly disorganized retinal structures.

### 5.3 Quantitative Sensitivity Gain

The improvement introduced by PFI was quantified by directly comparing the static and dynamic signal amplitudes (Fig. 5).

**Figure 5.**
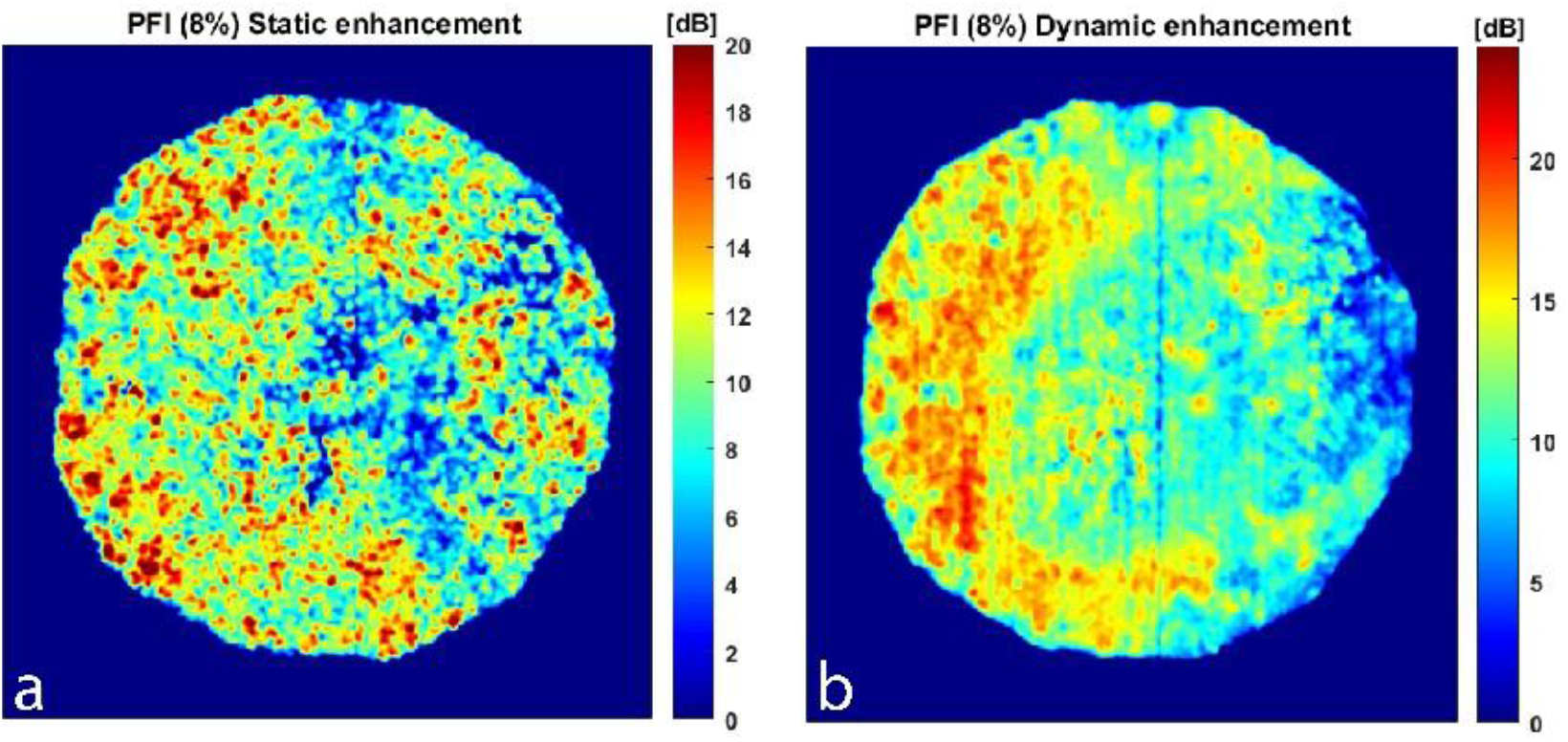
Quantification of signal enhancement produced by partial field illumination (PFI). **a** Static signal enhancement map calculated as the ratio of the static FFOCT signal obtained under partial field illumination (Fig. 4c) to that obtained under full-field illumination (Fig. 4a), displayed in decibels (dB). Both images were smoothed using a Gaussian filter prior to division. **b** Dynamic signal enhancement map calculated as the ratio of the Phase Fluctuation Index obtained under partial field illumination (Fig. 4f) to that obtained under full-field illumination (Fig. 4d), also displayed in dB after Gaussian smoothing. PFI produced a 19.8 dB improvement in static signal and a 23.2 dB improvement in dynamic signal, confirming that reducing incoherent background is particularly beneficial for dynamic functional imaging where useful signals are close to the detector noise floor.

Static signal enhancement was calculated from the ratio of the conventional 2-phase FFOCT images obtained under PFI and full-field illumination (Fig. 5a), while dynamic enhancement was quantified from the ratio of the corresponding Phase Fluctuation Index maps (Fig. 5b). Prior to division, both images were smoothed using a Gaussian filter with σ = 2 pixels to improve robustness of the comparison.

Quantitative analysis revealed a static signal improvement of 19.8 dB and a dynamic signal improvement of 23.2 dB when using 8% PFI compared with conventional full-field illumination.

The stronger gain observed in dynamic contrast confirms that PFI is particularly beneficial for D-FFOCT functional imaging, where the relevant biological features are often close to the detector noise floor. In this regime, reducing incoherent background is more effective than simply increasing total illumination power.

Importantly, these gains were obtained while preserving the same reference/sample ratio and without modifying the interferometric detection scheme, demonstrating that PFI acts as a direct sensitivity enhancement strategy rather than a change in image interpretation.

When combined with the ratio-free polarization architecture, PFI provides a cumulative sensitivity improvement of approximately 26.2 dB relative to conventional NPBS-based full-field D-FFOCT. This represents a major improvement for deep functional imaging of retinal organoids and establishes a practical route toward high-sensitivity label-free imaging of complex biological systems under photon-limited conditions.

## Discussion

This study demonstrates that sensitivity optimization in dynamic full-field optical coherence tomography (D-FFOCT) can be substantially improved through two complementary strategies: ratio-free optical flux redistribution and partial field illumination (PFI). Together, these approaches address the two principal limitations of deep dynamic imaging in highly scattering biological samples: inefficient use of optical power and detector saturation by incoherent background.

The ratio-free polarization architecture improves D-FFOCT sensitivity by enabling continuous tuning of the sample-to-reference field ratio while strongly suppressing parasitic reflections originating from conventional non-polarizing beam splitter (NPBS) configurations. Unlike standard 50/50 beam splitter systems, where the optical flux distribution is fixed and inherently inefficient, the present architecture allows direct adaptation to sample scattering conditions and imaging depth. This flexibility is particularly important in retinal organoids and other dense 3D biological models, where multiple scattering rapidly dominates the detector dynamic range and limits the detectability of weak dynamic signals.

Experimentally, this optical redistribution alone provides a 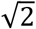-fold (3 dB) sensitivity improvement relative to conventional NPBS-based D-FFOCT. Although this gain may appear moderate compared with the larger improvements produced by PFI, it is especially valuable because it is achieved without increasing optical dose, modifying acquisition speed, or altering the biological preparation. It therefore represents a robust baseline improvement that can be directly implemented in existing systems with minimal modification.

Beyond sensitivity gain, the ratio-free architecture offers a major translational advantage for biomedical imaging. In retinal clinical applications, optical exposure must remain strictly below safety limits, and dose-limited operation is often the dominant practical constraint rather than source power availability. By decoupling sample illumination from reference intensity, the ratio-free configuration maximizes interferometric contrast while preserving strict biological irradiation limits. This is particularly relevant for future in vivo retinal imaging and for long-term monitoring of sensitive stem-cell-derived cultures.

When optical dose is not the primary constraint, the ratio-free architecture becomes even more advantageous because parasitic reflections are strongly suppressed, allowing substantially higher incident flux without saturation of the detector dynamic range. As shown in Fig.2a, under ideal source-power conditions and optimal ratio tuning, theoretical sensitivity improvement can exceed 25 dB.

Partial field illumination provides a second and substantially larger improvement by directly reducing incoherent background generated by multiple scattering. In highly scattering samples such as retinal organoids, this incoherent contribution often occupies a significant fraction of the camera dynamic range and prevents weak dynamic fluctuations from exceeding the system noise floor. The present results show that reducing the illuminated field dramatically improves both static and dynamic contrast, with a particularly strong effect on dynamic signal detection.

At 120 µm imaging depth, 8% PFI produced a 19.8 dB improvement in static FFOCT signal and a 23.2 dB improvement in dynamic D-FFOCT sensitivity. This stronger enhancement in dynamic contrast confirms that D-FFOCT is substantially more sensitive to incoherent background than conventional structural FFOCT. Functional imaging relies on weak temporal fluctuations rather than static strong reflectivity, and these fluctuations are often located near the detector noise floor. In this regime, reducing background contributions is more effective than increasing total illumination power.

The improved visualization of retinal rosettes further supports the hypothesis that scattering, rather than optical aberration alone, is the dominant factor limiting deep imaging performance in retinal organoids. Rosette structures are particularly difficult to image because local disruption of retinal organization increases both scattering and aberration. The strong improvement observed with PFI suggests that scattering mitigation should be considered a primary optimization target for future organoid imaging strategies.

Further reduction of the PFI ratio—potentially down to 0.5% as suggested by related studies—could provide even greater sensitivity gains and stronger evidence that scattering dominates over aberration in these systems. However, this improvement comes at the cost of increased acquisition time, since a smaller illuminated area requires proportionally more image tiles to reconstruct the same total field of view. In the present work, 8% PFI corresponds to approximately a 12.5-fold increase in acquisition time compared with full-field illumination.

While this trade-off may be acceptable for high-sensitivity applications such as photostimulation studies or detailed retinal organoid characterization, it may limit routine large-scale imaging workflows. Future implementations based on structured illumination, sparse illumination arrays, line scanning with descanned detection, or rolling-shutter synchronization could preserve the sensitivity benefits of PFI while reducing acquisition time penalties. In particular, sparse distributed illumination would likely be more efficient than the single-spot implementation used here for suppressing multiply scattered light.

The ratio-free architecture also establishes an optimal configuration for interface self-referenced D-FFOCT (iSR-D-FFOCT), where interference occurs between the sample and the substrate interface rather than an external reference mirror. This self-referenced design minimizes sensitivity to mechanical vibrations and phase instability while preserving high dynamic contrast. The improved detection of nuclei, membrane-associated structures, and nanometric interference fringes in Müller glial cells demonstrates the potential of this configuration for probing subtle intracellular dynamics and membrane fluctuations at very high sensitivity.

Such performance is particularly promising for future photostimulation experiments aimed at detecting osmotic responses, membrane deformation, and fine physiological changes associated with ion channel activation. Because these signals are often extremely weak and occur near the detection limit of conventional OCT systems, the combined ratio-free and PFI strategy provides a practical route toward functional imaging of previously inaccessible biological phenomena.

Overall, the cumulative sensitivity improvement of approximately 26.2 dB achieved in this work represents a major step forward for D-FFOCT applied to complex biological systems. More importantly, this improvement is obtained using experimentally accessible modifications that remain compatible with standard high-numerical-aperture FFOCT implementations. Rather than introducing a fundamentally new OCT modality, this work provides a practical framework for optimizing existing D-FFOCT systems for deeper, more sensitive, and more biologically relevant functional imaging.

These results expand the potential of D-FFOCT for retinal research, disease modelling, drug screening, and live-cell monitoring, and support its future translation toward clinical and translational biomedical imaging applications.

## Methods

### Optical setup

The ratio-free D-FFOCT system was built as a polarization-based Linnik interferometer (Fig. 1). Illumination was provided by a mounted light-emitting diode (LED; M810L5, Thorlabs, Newton, NJ, USA) centered at 810 nm with a spectral bandwidth of 30 nm, used as an extended spatially incoherent source (S1) with an emitting area of 1 mm^2^.

A first lens pair (L1; ACL25416U-B and AC254-150-B-ML, Thorlabs) imaged the source onto the illumination mask (OM). A first polarizer (Pol.1; LPNIRB100-MP2, Thorlabs), combined with a polarizing beam splitter (PBS; CCM1-PBS252/M, Thorlabs), controlled the optical flux ratio between the sample and reference arms by adjusting the balance between horizontal and vertical polarization components.

The illumination mask was relayed to the back focal planes of the two microscope objectives (Obj.1 and Obj.2; UMPLFLN20XW, Olympus/Evident, Japan) using a doublet (L2; AC508-150-B-ML, Thorlabs) and intermediate relay lenses (L3 and L4; AC508-150-B-ML, Thorlabs), forming the Linnik interferometer. A silver mirror (M1; PFE20-P01, Thorlabs) directed the beam into the interferometric arms.

The illumination iris (Ir.1), positioned at the focal plane of L2 and conjugated to the sample plane, was used for partial field illumination (PFI) to reduce incoherent background generated by multiple scattering. A second iris (Ir.2) was conjugated to the illumination path through the PBS to maintain optical alignment.

Dynamic and static phase shifts were introduced using a piezoelectric actuator (PZT; PK44M3B8P2, Thorlabs) and a linear translation stage (XR50P, Thorlabs), respectively. Zeroth-order quarter-wave plates (QWP1 and QWP2; WPQ10M-808, Thorlabs) rotated the polarization state after reflection from either the sample or the reference mirror, enabling polarization routing through the PBS and strong suppression of parasitic reflections.

A second polarizer (Pol.2) acted as an analyzer to project the orthogonal sample and reference fields onto a common polarization axis before detection by a cMOS camera (Q-2HFW, Adimec, Netherlands). A detection mask (DM) was positioned at the focal plane of the detection optics. Steering mirrors (M2, M4, and M5) were used for optical alignment, and the sample was mounted on a motorized 3D translation stage (RAMM RM-1250 and LS-100, ASI, USA).

### Simulation

The theoretical sensitivity analysis presented in Fig. 2 was implemented in MATLAB using the analytical expressions derived from the D-FFOCT signal model. Simulations were based on Eq. 2 and the NPBS reference sensitivity expression, using representative parameters corresponding to highly scattering biological samples: 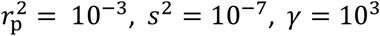. Where 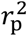represents the effective parasitic optical reflection term, *s*^2^ is the coherent sample reflectivity, and *γ* is the ratio between incoherent and coherent sample return.

These values were chosen to reproduce experimentally relevant imaging conditions encountered in deep retinal organoid imaging, where multiple scattering dominates the detector dynamic range. The influence of the polarization angles α and β on the ratio-free architecture was evaluated by calculating the theoretical signal-to-noise ratio (SNR) improvement relative to the conventional NPBS configuration.

The MATLAB simulation script used for generating the sensitivity maps and optimization curves is available on GitHub.

### Image Acquisition

Images were acquired using the ratio-free D-FFOCT system shown in Fig. 1. Illumination was provided by a mounted light-emitting diode (LED) source centered at 810 nm with a spectral bandwidth of 30 nm, corresponding to a coherence length of 9.62 µm.

The source was coupled to a water-immersion objective (UMPLFLN20XW, Olympus/Evident, Japan) with a numerical aperture of 0.5, resulting in a theoretical transverse resolution of 810 nm and an axial resolution of 3.65 µm [23].

All datasets were acquired over 5.12 s at 100 frames per second (FPS). The polarization angles α and β were adjusted experimentally to maximize SNR under these acquisition conditions. For standard D-FFOCT imaging, both angles were set to 10°. For interface self-referenced D-FFOCT (iSR-D-FFOCT), the optimal self-referenced configuration was obtained by setting α = 90° and β = 0°.

Partial field illumination (PFI) was implemented using the illumination iris (Ir.1 in Fig. 1). For PFI experiments, the illuminated area was reduced to 8% of the full-field configuration while maintaining the same reference-to-sample ratio. Exposure times were adjusted accordingly to preserve comparable detector filling conditions.

### Rolling-Phase Detection and Signal Demodulation

Dynamic data were acquired using the rolling-phase modulation regime previously described for D-FFOCT [26]. During acquisition, a continuous linear phase shift equivalent to 0 to 6π was introduced by controlled displacement of the piezoelectric mirror (PZT in Fig. 1) throughout the temporal sequence.

This rolling-phase detection avoids phase bias artifacts associated with static interferometric conditions and improves robustness against speckle and fringe artifacts for dynamic structures with limited drift.

Data were demodulated at 585 mHz to isolate the dynamic interferometric contribution and recover temporal fluctuations associated with intracellular activity.

For the PFI experiments, 512 raw full-field interferograms were acquired at 100 Hz for both full-field illumination and partial field illumination conditions.

### Dynamic Image Rendering

Three complementary metrics were calculated from the temporal fluctuations of the interferometric signal to characterize cellular dynamics.

The first metric was the recently introduced Phase Fluctuation Index (PFI), defined as: <|ΔI(t)|>, which emphasizes the magnitude of local phase fluctuations [26]. This metric is approximately linearly related to both scatterer reflectivity and the distribution of phase variability induced by intracellular active transport. Importantly, it is independent of the initial phase bias of the scatterers and therefore reduces artifacts such as speckle and stationary fringes.

A high PFI value may correspond either to a strong scatterer with limited motion or to a weak scatterer with large dynamic displacement.

Two additional metrics were computed from the magnitude spectral density (MSD): the mean frequency <MSD>, and its standard deviation StD(MSD) [24].

The mean frequency reflects the characteristic speed of intracellular transport and provides an estimate of cellular activity and motility, whereas the standard deviation helps distinguish directional transport from Brownian motion, thereby separating organized cellular motion from drifting extracellular material.

Together, these three metrics describe:

- dynamic magnitude through PFI,
- quantitative transport speed through mean MSD,
- motility versus Brownian behavior through MSD standard deviation.

These parameters were assigned to a Hue–Saturation–Brightness (HSB) color space for visualization:

- hue: mean MSD (<MSD>)
- saturation: standard deviation MSD (StD(MSD))
- brightness: Phase Fluctuation Index

In this representation, blue corresponds to slower intracellular dynamics, red to faster transport, low saturation to Brownian motion, and high saturation to directional transport.

Display thresholds were determined using percentile cutoffs of the signal distributions: hue: 0.1% to 99.9%, saturation: 5% to 99.9%, brightness: 5% to 99.9%.

### PFI quantification

In order to quantify the PFI gain in sensitivity (G_PFI_), the following coefficient was calculated:

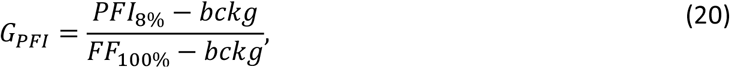

with PFI_8%_the pixel value of the PFI image at 8% (either 2-phase static or phase fluctuation index), bckg the background in each metric considered when the camera is shotnoise limited, and FF_100%_the pixel value of the standard Full-Field image.

### Sample Preparation

#### Retinal Organoids

Human retinal organoids at day 150 of differentiation were maintained in 12-well plastic-bottom culture plates (Corning® Costar® Not Treated Multiple Well Plates). Before imaging, each well was filled with pre-warmed fresh culture medium.

The organoids were kept in the dark for at least 1 h in a humidified incubator at 37°C and 5% CO(_2) prior to imaging.

### Müller Glial Cells

Human induced pluripotent stem cell-derived Müller glial cells (hiMGC; PMID: 33683746) were cultured in glass-bottom dishes precoated with Geltrex®.

These cultures were used for interface self-referenced D-FFOCT (iSR-D-FFOCT) imaging to evaluate fine intracellular dynamics and membrane-associated structures under high-sensitivity self-referenced conditions.

## Data availability

Source data for the graphs in the main figures is available as Supplementary Data, and any remaining information can be obtained from the corresponding author upon reasonable request.

## Code availability

MATLAB scripts used for sensitivity simulations are available at: https://github.com/Tual29/ratio-free-DFFOCT/tree/main

## Acknowledgements

We thank Camille Hourton and Sacha Reichman for the sample preparation of the Müller glial cells, Olivier Goureau and Teo Bendahmane for providing retinal organoids.

We are very grateful to Olivier Thouvenin for fruitful exchange on partial field illumination, and Kate Grieve for supporting our research.

The author wishes to acknowledge funding from IHU FOReSIGHT “RETINA-LENS” [P-RETION-IHU-000] and ANR “No-D-EYE” [M25JRAR029].

## Contributions

T.M. conceived the study, designed and built the optical setup, performed the theoretical modelling and numerical simulations, carried out the experiments, acquired and analyzed the data, interpreted the results, and wrote the original manuscript.

